# An Adverse Outcome Pathway for Food Nanomaterial-induced Intestinal Barrier Disruption

**DOI:** 10.1101/2024.10.11.617731

**Authors:** Deborah Stanco, Dorelia Lipsa, Alessia Bogni, Susanne Bremer-Hoffmann, Laure-Alix Clerbaux

## Abstract

**Introduction:** Ingestion of nanomaterials (NMs) might impair intestinal barrier, but the underlying mechanisms remain evasive, and evidence is not systematically gathered or produced. A mechanistic-based approach would be instrumental to assess if relevant NMs disrupt intestinal barrier to support NM risk assessment in the food sector.

**Methods:** Here, we developed an adverse outcome pathway (AOP) based on biological plausibility and by leveraging existing information of an existing NM relevant AOP leading to hepatic outcomes. We then extracted the current evidence existing in the literature for a targeted selection of NMs with high food sector relevance, namely ZnO, CuO, FeO, SiO_2_, Ag NMs and nanocellulose.

**Results:** We propose a new AOP (AOP530) that starts with endocytic lysosomal uptake leading to lysosomal disruption inducing mitochondrial dysfunction. Mitochondrial impairments can lead to cell injury/death and disrupt the intestinal barrier. The evidence collected supports that those food NMs can be taken up by intestinal cells and indicates that intestinal barrier disruption by Ag, CuO, SiO_2_ NMs might occur whilst only few studies support that outcome for FeO, ZnO. Lysosomal disruption and mitochondrial dysfunction are rarely evaluated. For nanocellulose, none of the studies report toxic-related events.

**Conclusions:** Collecting the existing scientific evidence supporting our AOP linking NM uptake to intestinal barrier impairments allowed us to highlight current evidence gaps but also data inconsistencies. Those latter could be associated with the variety of stressors, biological systems and KE-related assays used in the different studies, calling for further harmonized methodologies and production of mechanistic evidence in the safety regulatory assessment of NMs in the food sector.

## 1. Introduction

### 1.1 Regulatory needs in risk assessment of food nanomaterials

Nanotechnology presents many possibilities for the food industry, offering potential benefits in areas like targeted nutrient delivery, improved food preservation, and enhanced sensory experiences. However, due to their unique properties at the nanoscale, nanomaterials (NMs) might necessitate additional testing requirements. To implement the information requirements, the European Food Safety Authority (EFSA) has published two guidance documents outlining criteria for evaluating materials at nanoscale. These guidance documents address defined “engineered nanomaterials” (1) as well as food additives with a fraction on smaller particles (2). The EFSA guidance recommends a tiered analysis of nanospecific considerations revolving around the behavior of the NM within the gastrointestinal (GI) tract. This includes dissolution dynamics, cellular uptake, transcytosis, and potential disruption of the intestinal barrier. Additionally, genotoxicity and the accumulation potential of the NM are assessed. This tiered analysis offers a unique opportunity to integrate *in vitro* testing into the regulatory process. The guidance document proposed additionally a stepwise approach, focusing on *in vitro* assays related to cytotoxicity (cell death), oxidative stress, (pro)inflammation, and impairment of the intestinal barrier. Furthermore, *in vitro* dissolution testing under simulated lysosomal and GI conditions, are recommended to obtain a more exhaustive picture on the complex biological interactions of ingested NMs in the GI tract In order to develop case studies implementing the two guidance documents, EFSA launched a pilot project using nanocellulose (NC) as an emerging material in the food sector and additional case studies are ongoing in the remit of an EFSA funded project called NAMS4NANO where various NM are characterized and tested in different *in vitro* models (3). The current evidence on a potential intestinal barrier disruption triggered by those NMs relevant in the food sector has never been collected within a mechanistic proposed approach so far.

### 1.2 Intestinal barrier disruption, an adverse outcome triggered by food NMs?

The gut barrier plays crucial roles by acting as a protective barrier that separates the internal blood from the luminal content but also as a selective barrier that regulates fluxes of water, ions and essential dietary nutrients. The intestinal barrier consists of a multilayer system encompassing a chemical layer containing the antibacterial proteins secreted by Paneth cells, a mucus layer secreted by goblet cells, an epithelial layer and the cellular immune system. Enterocytes and goblet cells are the most studied cell types. Alteration of one or many of those layers leads to increased intestinal permeability, also called intestinal hyper-permeability or leaky gut syndrome, enhancing translocation of bacteria, bacterial products (such as lipopolysaccharides) and undigested nutrients from the intestinal lumen into the systemic circulation (4). Many diseases arise or are exacerbated by a leaky gut, including diarrhea, inflammatory bowel disease, celiac disease, autoimmune hepatitis or type 1 diabetes (5–8). In addition, individuals with a leaky gut are more vulnerable to toxicity driven by chemical exposure, as dietary pesticides or food chemicals can more easily enter the systemic blood and reach the target organs (9).While it can be argued that intestinal barrier disruption is an intermediate event towards various clinical outcomes, in view of its central role, we advocate here to consider this event as an adverse outcome when assessing the toxicity of food NMs.

### 1.3 Development of an Adverse Outcome Pathway (AOP) to support NM risk assessment in the food sector

While the underlying mechanisms remain evasive and evidence not systematically gathered or produced, a mechanistic-based approach would be highly instrumental to assess the impact of food NMs on intestinal barrier integrity. Such a risk assessment based on mechanistic reasoning requires relevant AOPs. The OECD defines the AOP as a description of a logical sequence of causally linked events (Key Events, KEs) at different levels of biological process, which follows exposure to a stressor and leads to an adverse outcome (AO) in humans or wildlife (10). Hence, a KE describes a measurable and essential change in a biological system that can be quantified in experimental or clinical settings. Then the strength of the relationship between two KEs (Key Event Relationship, KER) is established first by biological plausibility (e.g. binding to a receptor induces a signaling cascade) and second by weighting the current evidence supporting this causal link which then can highlight gaps in the knowledge or inconsistencies between studies. Data discrepancies can be due to different experimental settings but also to physicochemical properties in the case of NMs. The development of an AOP is a dynamic process building from hypothesized KEs towards refinement of KEs and KERs by assembling evidence found in literature or newly produced (11). An AOP development guidance was established by the OECD (12) and already developed AOPs describing various AOs are stored in the publicly accessible repository platform, called AOP-Wiki (https://aopkb.oecd.org/). There is no explicit quality assurance for AOPs in the AOP-Wiki, except when they have gone through the review process of the OECD (OECD 2018), making those AOPs compliant for potential regulatory use.

Initially developed to support new approach methodologies (NAM)-based chemical risk assessment, the framework was recognized as instrumental for prioritizing and developing testing strategies for NMs (13). Most AOPs developed for chemicals should be also applicable to NMs, considering certain adaptations at the MIE and early KEs levels (14). For example, AOP144 leading to liver fibrosis was proven to be nano-relevant as endocytic lysosomal uptake is described as a nano-relevant MIE connected to lysosomal disruption and mitochondrial dysfunction leading to cell death in the liver (15–17). Simple *in vitro* models for testing the potency of NMs were associated with the MIE and early lysosomal and mitochondrial KEs in this AOP (16). These early KEs lead to cell death.

Building on “tissue injury”, Halappanavar et al proposed a methodology for identifying NM relevant KEs based on plausibility, measurability and regulatory importance of the KEs (13). The authors identified inflammation (increased pro-inflammatory mediators, leukocyte recruitment/activation), oxidative stress and cell death as main upstream KEs to tissue injury. Reported endpoints and associated assays were identified allowing for quantifiable measurement of these KEs. Finally, in line with the increasing recognition of the central role of leak gut, the ‘intestinal barrier disruption’ was recently added in the AOP-Wiki in the COVID-19 context (14). But, even if food NMs are ingested raising the possibility of an adverse outcome on the intestinal barrier, there are currently no AOPs supporting the toxicity on the intestinal barrier following intestinal cellular uptake of ingested NMs.

In this study, we developed an AOP proposing a toxicological pathway towards intestinal barrier disruption following intestinal uptake of NMs via lysosomal and mitochondrial dysfunction. This AOP was developed by leveraging existing information of existing AOPs published in AOP-wiki and based on biological plausibility. Following the establishment of the AOP, we extracted current evidence in the scientific literature for a targeted selection of NMs with high food sector relevance. These NMs are ZnO, CuO, FeO, SiO2, Ag NMs and nanocellulose. Our in-depth literature review allowed us to highlight current knowledge gaps and data inconsistencies guiding future research.

## 2. Methods

### 2.1 AOP-wiki search to identify intestinal AOPs with NMs reported as stressors

In the prototypical stressors section of the AOP-Wiki, the keywords “nanoparticles”, “NPs”, “nanosized particles”, “nanomaterials”, “nanotubes” were used to filter AOPs in the AOP-Wiki which report a nanosized material as a stressor. For each AOP, the AOP number, MIE, AO and status information have been extracted.

### 2.2 AOP development strategy

The AOP was built first based on biological plausibility and by leveraging existing KEs in another NM relevant AOP. We started from the apical outcome ‘intestinal barrier, disruption’ that we postulated was due to ‘cell death/injury’ of the different intestinal cell types, identified as a main KE upstream of tissue injury in nano relevant AOPs (13) and with the rationale that intestinal barrier is composed of different layers of specific cell types, which death or injury impairs the barrier function. In this study, we started by focusing on enterocytes and goblet cells. The identification of the intermediate cell specific KEs between ‘cell death/injury’ and ‘intestinal barrier disruption’ was based on biological plausibility and was further refined following the literature search. On the other side of the pathway, the MIEs and initial KEs defined in nano relevant AOP in liver considering NM-induced ‘mitochondrial dysfunction’ leading to ‘cell death/injury’ were extrapolated by a tissue analogy-based approach as upstream KEs (Figure 1).

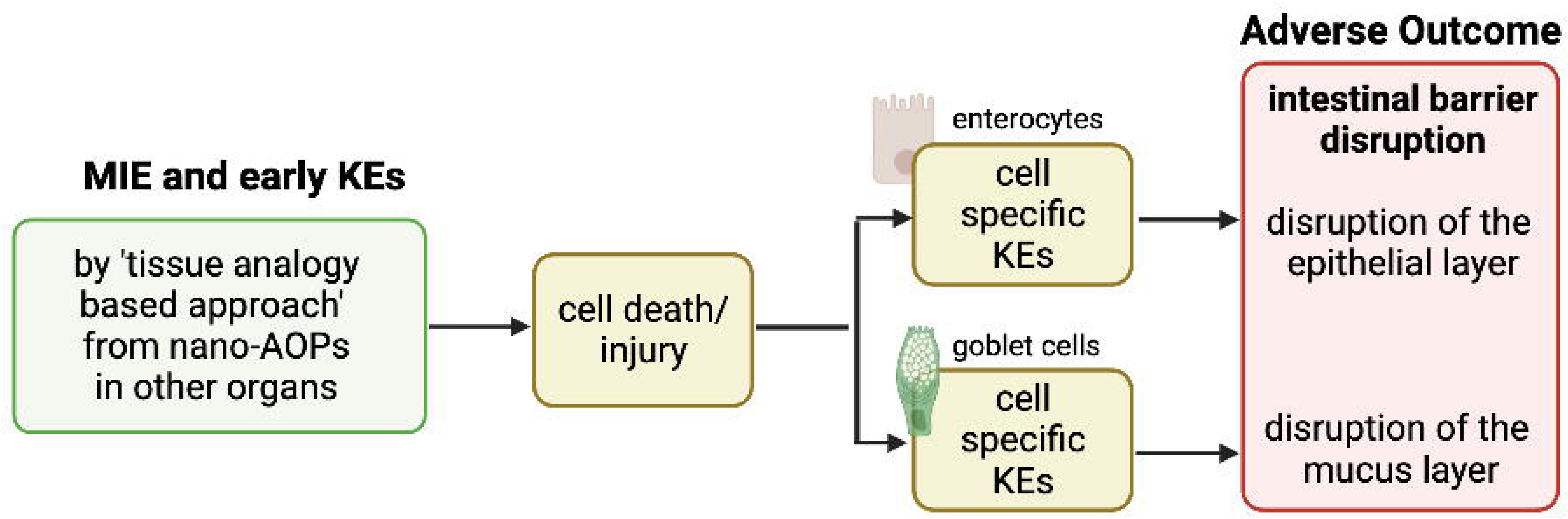

To verify the relevance of the proposed AOP for the food sector, we focused the literature research on NMs used as food additives and novel foods. Zinc oxide (ZnO) NM is a micronutrient supplement and food fortification ingredient can release Zn ions intracellularly after being absorbed by enterocytes (18). Silicon dioxide (SiO_2_) NM (E551) is a food additive that exhibits size-dependent cytotoxicity in *in vitro* studies (19). Iron oxides (FeO) NM (E17) serve as food additives and flavorings (20), the impact on the intestinal barrier is yet to be explored. The nanosilver (Ag NM, E174) is an antimicrobial material used as a colorant or food contact material that most properly exert its toxicity through silver ion release, but conclusive studies are lacking (21). Copper oxide (CuO) NMs are used as nutrients, food supplements, and pesticides and share potential ion-driven toxicity mechanisms with ZnO and Ag NMs (22).

### 2.3 Literature search

The literature search was performed both automatically using AOP helpfinder and manually in PubMed based on selected keywords, followed by a selection procedure of the relevant studies based on predefined exclusion criteria.

#### 2.3.1 Keywords selection and literature screening

The automatic search was performed using the AOP-helpFinder tool (23), a hybrid approach that combines text mining procedure and graph theory to automatically identify literature containing co-occurrence between the stressor(s) and KE(s) of interest. To do so, two lists of keywords have created based on expert knowledge:

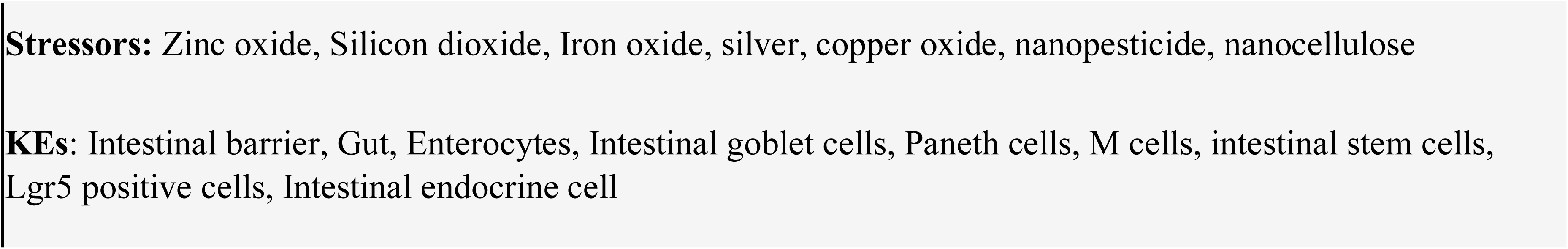

The parameters used to screen the abstracts in AOP helpfinder were (i) dismiss the first 20% of the abstract and (ii) apply the refinement filter, a second analysis performed using a lemmatization process which contextualized words with common stems (e.g. tests, testis, test). The abstracts were provided as outputs in a table along with PMID identifiers.

Then the titles and abstracts of publications within the PubMed database (as of March 2024) was screened using the keywords:

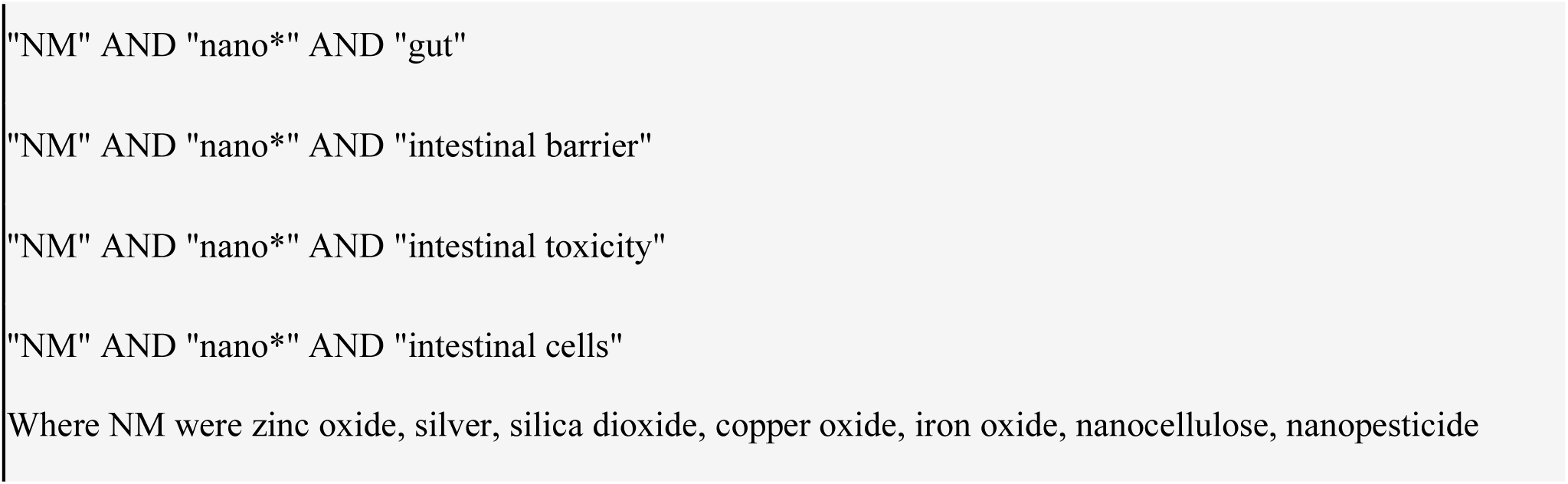

The abstracts were provided as outputs in a table along with PMID identifiers.

##### Exclusion criteria

For both literature search, we then manually excluded abstracts that were in duplicate, not in English, not containing primary data (e.g. reviews), not related to food NMs, not related to intestinal barrier or related to nanomedical applications. In addition, studies with human cells and samples as well as rodent studies were selected while studies using environmentally relevant species (e.g. worms, flies. Daphnia, fish) were excluded.

### 2.4. Extraction of relevant information

#### Collecting evidence

The entire publication of the identified abstracts was read by the authors. First, the origin of the NM was reported followed by the biological system used in the study (e.g. intestinal human cells or rodent studies). Then for each KE, the information regarding the measure was reported as *equal, increased or decreased* compared to control (e.g. experimental setting without exposure to the NM). The lowest dose at which a change was observed was retrieved. When no significant differences were observed, the highest dose used in the study was recorded. Only when the duration of exposition was different from 24 hours, the information was reported. In addition, the type of assays used to produce the evidence was retrieved from the studies. Finally, the references of the publication were collected.

#### Physicochemical score

To evaluate the level of characterization reported by the authors in the studies we allocated 0 point is nothing was mentioned regarding the size, 0.5 point if size of pristine was measured and 1 point if both pristine and NMs size in experimental medium was measured. If one other property was described, we allocated one extra point. Finally, if more than one property in extra to size measurement was measured, we allocated two extra points. This arbitrarily assigned grades ranging from the lowest (0) to the highest (3) number of NM measurements performed. Of note, for compounds coming from the JRC repository (24,25), the highest grade of 3 was given as they are thoroughly characterized.

## 3. Results

### 3.1 AOPs reporting NMs as stressors in the AOP-Wiki

Sta. The AOP144, currently under revision by OECD, refers to the correlation between the NM cellular uptake by endocytic lysosomal absorption, and the consequent mitochondrial dysfunction toward liver fibrosis (17). Others AOPs illustrate the NM-mediated lung toxicity (shown as grey in Table 1). AOP237, 303, 302, and 319 are under development as part of different OECD projects. Finally, there are several AOPs proposed that are not currently part of the OECD process. Not pertaining to the lungs, AOP209 connects silica nanoparticle-induced disruption of cholesterol to hepatotoxicity, while AOP207, AOP208, and AOP210 suggest pathways leading to infertility.

In addition, outside the AOP-Wiki, at least five AOPs identifying NMs as stressors and leading to lung outcomes are described in the literature in humans. Two AOPs starting from increased substance interaction and leading to lung emphysema and lung fibrosis, respectively have been proposed (26). Literature reports suggesting potential hazards of TiO_2_ served for the development of one putative AOP leading to lung cancer (27,28). AOP for graphene-family nanomaterial-induced lung damage was developed (29) as well as the AOP describing Ag NM toxicity towards the respiratory tract (30). Regarding intestinal outcomes, plausible AOP was proposed following ingestion of TiO_2_ nanoparticles which eventually lead to colorectal cancer (31). This AOP is not included in the AOP-Wiki. An AOP-oriented study also assessed cytotoxicity, oxidative stress, genotoxicity, perturbation of cell cycle and apoptosis on human intestinal cells upon Ag NMs exposure based on KEs present in AOPs reporting Ag NMs as a stressor but no intestinal outcomes have been reported (32).

Thus, even though food NMs enter the human body through oral route raising possibility of adverse effects on the gut barrier, there are currently no AOPs developed supporting the toxicity on the intestinal barrier function following cellular uptake by intestinal cells of ingested NPs. A structured approach to assess these effects can be instrumental. Here, we aimed to develop an AOP specifically focused on the impact of NMs on the intestinal barrier. This AOP construction relied on biological plausibility to link NM cellular uptake with disruption of the intestinal barrier. We then sought to evaluate this framework by extracting evidence from published research. Finally, our analysis aimed to identify any data gaps or inconsistencies in the evidence supporting this AOP.

### 3.2 Biological plausibility for an AOP linking NM uptake to intestinal barrier disruption

NM-relevant MIEs identified in AOPs in other tissues might be linked to the intestinal tissue. AOP144 is at the second highest stage of AOP development. The MIE *endocytic lysosomal uptake* leads to *lysosomal disruption*, which induces *mitochondrial dysfunction* leading to *cell injury/death*. This is of interest as recently a proposed AOP network linked NM-induced mitochondrial dysfunction to existing AOs in lung, liver, cardiovascular and nervous systems (Murugadoss et al., 2023). The authors identified that NM-induced mitochondrial toxicity is crucial for many tissues but interestingly did not mention the intestinal epithelium.

Hence, we propose an AOP that is biologically plausible (Figure 2).

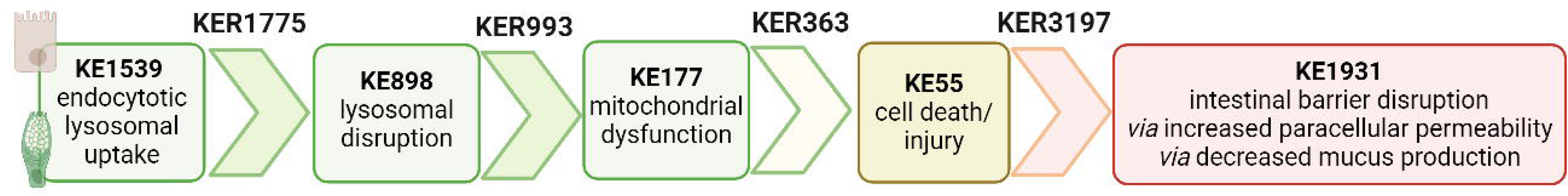

#### 3.2.1 NM endocytosis to intestinal cell death/injury

The biological plausibility that endocytic lysosomal uptake of NMs leads to lysosomal disruption is high and described in **[KER1775](https://aopwiki.org/relationships/1775)** in the AOP-Wiki. Briefly, endocytosis, discovered by Christian de Duve, is an active transport in which molecules are transported into the cell by engulfing them in plasma membrane, which then form a vesicle containing the ingested material inside the cell. Vesicles rapidly fuse to form larger compartments, known as endosomes. As detailed in KE1539 in the AOP-Wiki, there are different assays to evaluate NM cellular uptake (Box 1). Internalized material by endocytosis is then transferred to lysosomes, vacuoles containing hydrolytic enzymes in an acid milieu to degrade ingested material (33). Regarding NMs, once they are taken up by a cell and transported to the lysosome, the acidic milieu herein can either enhance their solubility, or they remain in the initial nano-form. Both situations can cause lysosomal swelling, followed by lysosomal disruption and the release of pro-apoptotic proteins (34,35). Lysosome disruption can be measured as in Box 2 from KE898 in AOP-Wiki.

The biological and causal link between lysosome and mitochondrial toxicity is detailed in **[KER993](https://aopwiki.org/relationships/993)**. Release of lysosomal proteases due to lysosomal disruption induces mitochondrial dysfunction (KE177) which encompasses a wide variety of changes in the structure and function of the mitochondria. The most reported ones are altered production of ATP, loss of mitochondrial membrane potential (MMP), inhibition of protein complexes in the electron transport chain, failure to produce enzymes that detoxify ROS (Box 3).

Finally, the biological plausibility of mitochondrial toxicity leading to cell death/injury is well established in the literature and captured within **[KER363](https://aopwiki.org/relationships/363)** included in OECD-endorsed AOP48 and in AOP144 under review in the AOP-Wiki (36). Cytotoxicity (apoptosis/necrosis) can be measured via different assays as detailed in Box 4 from KE55.

#### 3.2.1 Intestinal cell death leads to intestinal barrier disruption

The biological plausibility that damaged/dying enterocytes or goblet cells leads to increase of paracellular permeability due to disruption of the epithelial monolayer and to decrease of mucus secretion and thickness, respectively, is high but the causal link is not captured in the AOP-Wiki. We created **[KER3197](https://aopwiki.org/relationships/3197)** in the AOP-Wiki linking cell death/injury to intestinal barrier disruption. The intestinal barrier is a multilayer system composed of a chemical layer containing the antibacterial proteins secreted by Paneth cells, a mucus layer secreted by goblet cells, a one-cell-thick epithelial layer attached together through tight junction (TJ) proteins as well as a cellular immune layer. Intestinal permeability describes the movement of molecules from the lumen to the blood, and as such, is the measurable feature of the intestinal barrier function. Transcellular permeability encompasses passive diffusion from the apical to the basal side (lumen to blood), vesicle-mediated transcytosis and membrane receptor-mediated uptake. Paracellular permeability is regulated by the tight junctions between adjacent cells and by the integrity of the epithelium. The disruption of the intestinal barrier (KE1931) can be caused by damaging one or many layers due to cell death/injury of the specific associated cells. In this study, we will focus on enterocytes and goblet cells and associated alteration of epithelial monolayer integrity and mucus layer, respectively. Paracellular permeability and mucus secretion/thickness can be measured as described in Box 4.

### 3.3 Evidence Assessment

#### 3.3.1 Selection of relevant publications

Based on the keywords selected, we obtained 1249 papers with the combined Pubmed and AOP helpfinder search. We excluded 140 for ZnO, 137 for SiO_2_, 77 for FeO, 588 for Ag, 143 for CuO and 14 for NC papers because there were replicates or overlapped between the two-method searches, not in English or because the studies did not focus on NMs effect on intestinal barrier in humans or rodents (Figure 3). For nanopesticides, no results were obtained from either search.

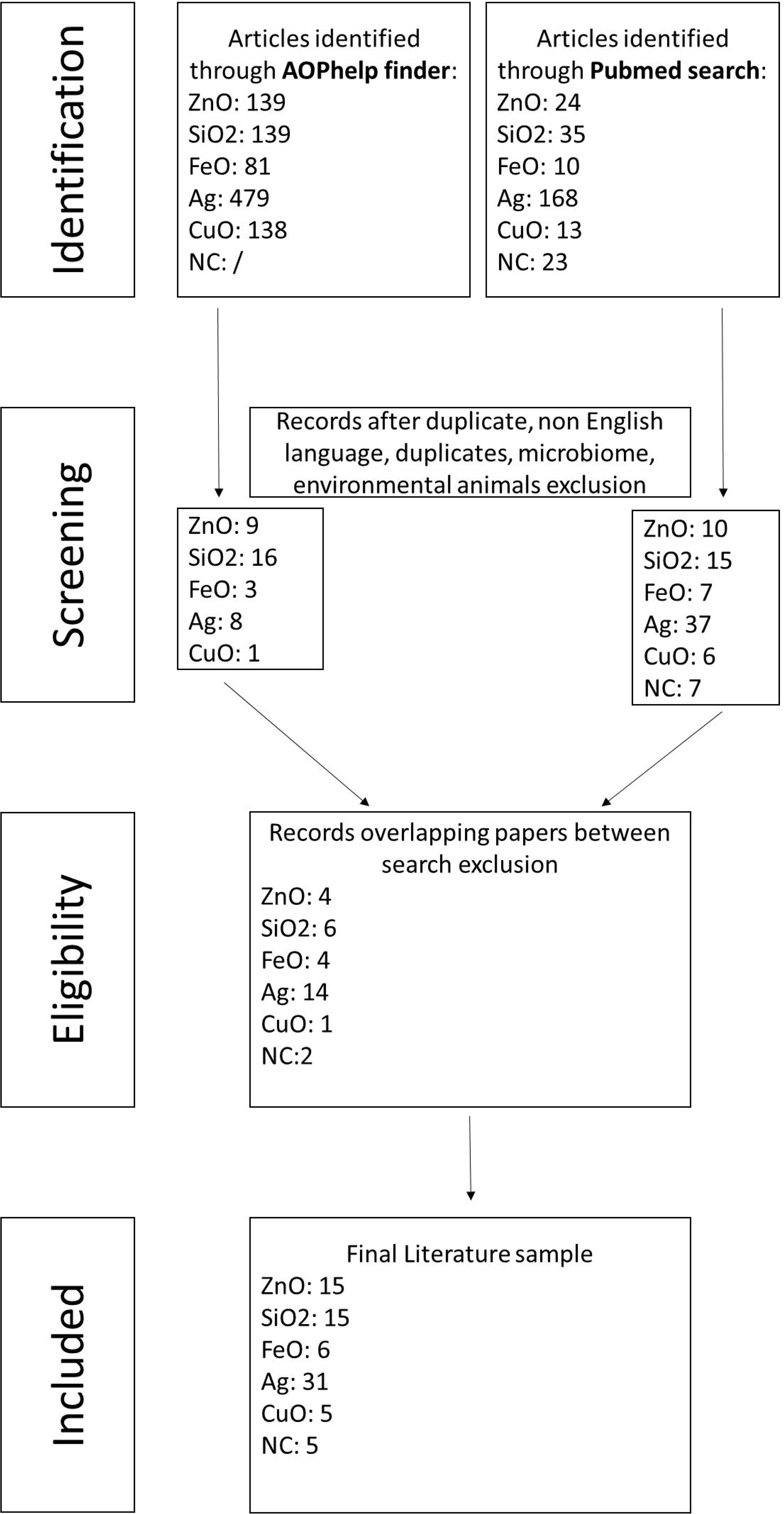

#### 3.3.2 Empirical support for KERs

For each selected publication, we reported the stressor used in the study along with a physicochemical score (PC). In their guidance, EFSA insists on the importance of a precise physicochemical characterization of NMs by reporting composition, size, size distribution, shape, charge, agglomeration state, surface composition. The level of NM characterization in the studies were reported as grades ranging from the lowest (0) to the highest (3) number of physicochemical properties measured. Then we reported the biological system used, including cell type, proliferating (P) or differentiated (D), cultured in transwell (T) or glass side (G) and if the treatment was applied at the apical (AP) or basolateral (BL) side. For rodent studies, the route of exposure, duration and dose were retrieved. Then the measure regarding each KE was reported as equal (=), increased (+) or decreased (−) compared to control. For example, if TEER values decreased upon exposure to NM, this means the intestinal barrier disruption increases (+). The dose indicated is the lowest inducing a change (+/−) in the study and the highest dose used in the study when no changes were reported (=). The type of assays used to produce the evidence was retrieved from the studies (Tables 2-7).

**[MIE](https://aopwiki.org/events/1539).** Consistent data for ZnO, CuO, FeO, SiO_2_ and Ag NM are recorded for *endocytic lysosomal uptake*, by using different imaging techniques (i.e CLSM, SEM, TEM) and ICP-MS, -OES or AES, for visualization and quantification, respectively (Tables 2-6). NC internalization has not been investigated so far (Table 7), probably because of its challenging organic carbon-based nature that requires specific immunofluorescence staining for detection (3). In a previous study, we stained CNC, NFC and BNC by Calcofluor and CBM-GFP and the presence of NM inside the cells was analyzed by measuring the emitted fluorescence by CLSM.

**[KER1775](https://aopwiki.org/relationships/1775) and [KER993](https://aopwiki.org/relationships/993).** Very few data evaluated *lysosomal disruption* after NM uptake in the gut. They are primarily related to Ag NMs assessed by using proteomics techniques (Table 6) without being confirmed with other specific assays such as lysosomal staining or specific lysosomal disruption assays. Hence the weight of evidence for KER1775 is low. Similarly, as lysosomal disruption is almost never assessed, evidence linking it to mitochondrial disruption is almost inexistent, hence empirical evidence is low for KER993.

**[KER363](https://aopwiki.org/relationships/363).** Few evidence supports *mitochondrial disruption* obtained by no specific assays across CuO, SiO_2_, and NC studies (Table 3,5, 7). The *cell death/injury* was investigated via several assays such as MTT, LDH release, CellTiter-Blue, and DAPI staining *in vitro* after NM exposure. Studies using ZnO NMs supports a causal relationship between mitochondrial dysfunction and cell death, as assessed by MTT, MMP, WST-1 or TEM (Table 2). However, six studies reported both mitochondrial dysfunction and cytotoxicity while four other studies using the same ZnO NMs showed consistently cellular uptake without mitochondrial and cellular damage (Table 2) (51–55). Several studies indicated a cytotoxic effect of CuO NMs (Table 3). While assessed in more than half of the selected studies, minor or no significant cell deaths or injuries were reported for SiO_2_ (Table 5) (46,47), while none evaluated mitochondrial damage (Table 5). Regarding Ag NM, several reports indicated changes in mitochondrial membrane potential (Table 6). In particular, smaller silver nanoparticles with higher surface activity led to notable mitochondrial changes and increased oxidative stress (44,45). Similar observations are reported for cell death which is influenced by size, coating, and surface charge of Ag NMs (49,50) (Table 6). Regarding NC, only one publication reported both mitochondrial and cytotoxicity dysfunction of the differentiated Caco2 monolayer after crystal (CNC) and fibrillar (FNC) nanocellulose exposure (48). Interestingly, among the four types of CNC tested with several nano-scale dimensions, only the one with the medium size induced a decrease of mitochondrial functionality and cell viability. Moreover, among the three FNC tested, the one with the intermediate dimension (80nm) caused mitochondrial dysfunction. Based on those data, empirical evidence for KER363 is considered as moderate for food NMs.

**[KER3197](https://aopwiki.org/relationships/3197).** Temporal and dose-response evidence to support that enterocytes or goblet *cell death/injury disrupts intestinal barrier* function is limited with some inconsistencies across studies. For ZnO NMs, seven studies investigated both cell injury and intestinal barrier of which four studies using the same NMs observed no changes in both events *in vitro*, while in another *in vitro* study, digested ZnO NM increased intestinal barrier. One mouse study noted that gut mucosa was disrupted following intragastric administration of ZnO NMs with decreased of *Cldn3* mRNA expression, suggesting impairment of intestinal barrier integrity while another mouse study further supports damaged intestinal barrier but without histological cell damage (ref). Regarding CuO NMs, Ude et al observed uptake, cell death and barrier disruption supporting that CuO NM might trigger differentiated Caco2 to death-induced barrier disruption (Table 3) (56). In a subsequent study, they confirmed that outcome in Caco2/RajiB and Caco2/RajiB-MTX barrier *in vitro* models by using the same type of CuO NM (57). Besides, morphological changes of microvilli were also reported by SEM images supporting the results related to barrier disruption. Interestingly, Li et al showed that differentiated Caco-2 taking up CuO NM possessed morphological ultrastructure changes (58). The effect on the organelle’s morphology could be related to cell death. Indeed, the increased number of vacuoles observed in mitochondria within the cells may be attributed to the escape of NMs from the endocytic pathway. Bypassing lysosomal degradation, these NMs are released into the cytoplasm, leading to impairments in organelle structure and function (59). Regarding FeO NMs, various reports showed cytotoxic effects without causing intestinal barrier dysfunction *in vitro* (Table 4). Long-term oral administration of nano-iron oxide in mice caused intestinal damage with loss of villi structures and hepatic dysfunction (60). Upon ingestion of SiO_2_ NM, one mouse study observed cell and intestinal damage, while another did not observe any changes in rat fed with Kaolinite (Table 5). Three *in vitro* studies reported no cell death nor intestinal damage while two others pointed towards intestinal barrier disruption without evidence for cellular damage (ref). Ag NM literature is the most prolific in assessing cell and intestinal damage, with 24 studies supporting cell death/injuries and six showing no cytotoxicity (Table 6). A causal relationship toward intestinal barrier disruption is supported by four studies while six others observed cell death without gut barrier disruption. Regarding NC, no negative effect on the intestinal epithelium has been reported in the majority of the studies (Table 7). Hence, empirical support is proposed as moderate for KER3197.

**[AO](https://aopwiki.org/events/1931).** The *intestinal barrier disruption* is commonly monitored by assessing transepithelial resistance (TEER), TJ expression profile (qRT-PCR, IF) or by measuring the apparent permeability coefficient (Papp) of the membrane generally with Luciferase yellow or FITC-dextran assays. While intestinal barrier disruption is partially reported across the studies, the results are heterogeneous in some cases. Barrier disruption was observed *in vitro* and *in vivo* upon SiO_2_ NMs exposure but not consistently (Table 5). *In vitro* exposure to Ag NMs led to decreased TEER values, changes in gene expression of TJs and cellular impedance, supporting impairment of the barrier integrity (61–67). Moreover contradictory Ag NM dose-response data have been observed with both low and high doses causing barrier impairment (49,62). Importantly, this variability may depend on the experimental conditions, both the *in vitro* system and nanoparticles used in the studies. For instance, the intestinal barrier disruption assessment by TEER was reported only in some of the studies whereas others based their results by indirect observation of cytotoxicity and other criteria such as genotoxicity and proteomics (65,68–70). Finally, no significant barrier disruption in NC studies has been observed so far (Table 7).

This AOP is mostly qualitative as empirical support for the dose concordance is not well established for all of the KERs in the pathway. Additional studies are needed to support the essentiality of the KES and to provide evidence on temporal and dose-response relationships for each KER. Based on this data extraction, we summarized the upstream and downstream KEs and weight-of-evidence evaluation of KERs for NMs relevant for the food sector in Table 8.

**Table 8.**
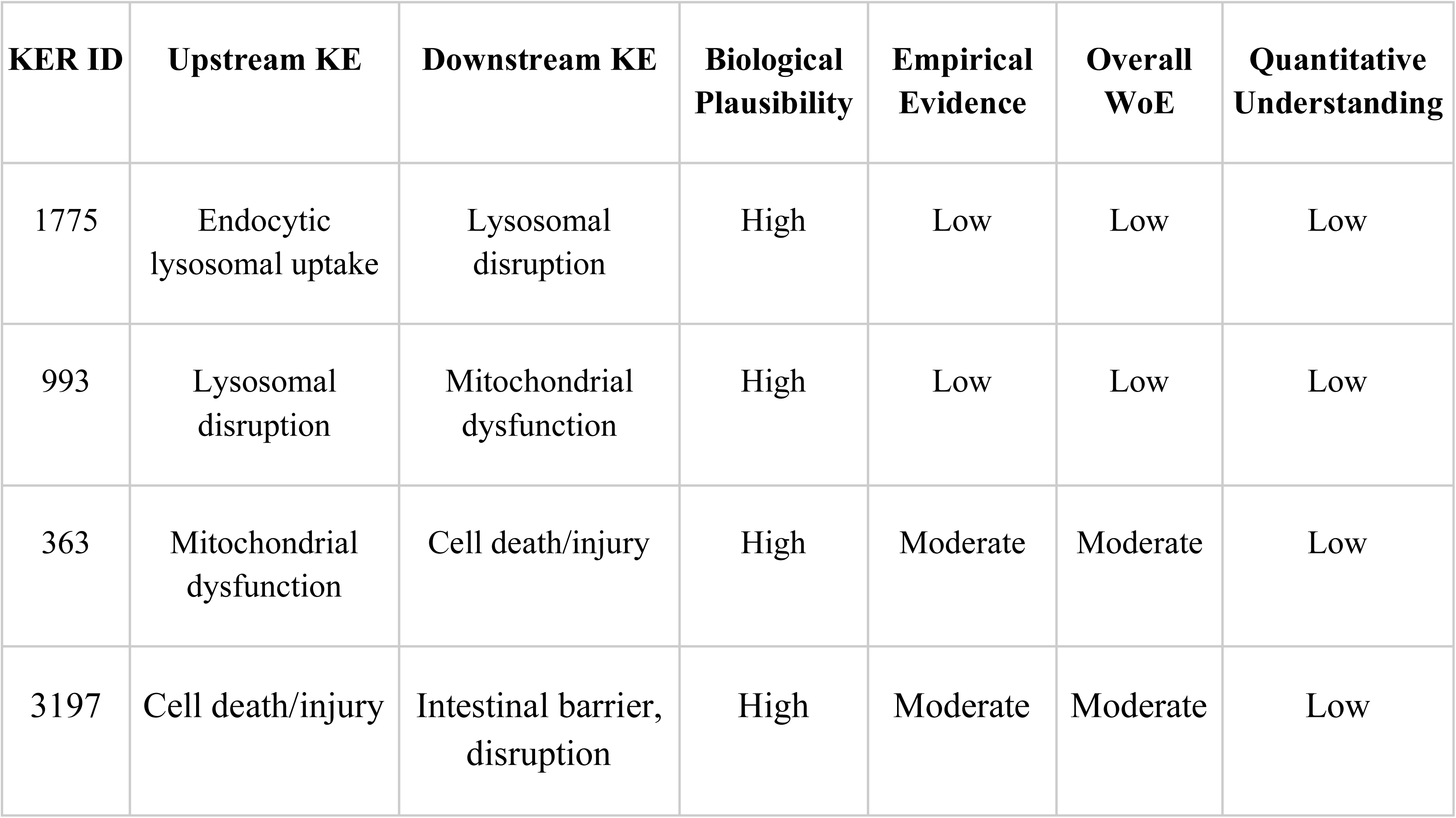
Summary of weight-of-evidence (WoE) evaluation of KERs in the AOP.

### 3.4 Uncertainties, inconsistencies and critical gaps

The above table highlights evidence gaps, particularly for KER1775 and KER993 as lysosomal disruption was almost never evaluated (Table 8). Besides, our quantitative understanding is currently limited. In addition, data were not consistent among studies that used various stressors, biological systems and E-related assays.

#### Stressors

In general, establishing a clear link between the physicochemical properties of NMs and their uptake into cells is challenging due to the complex interplay of properties. Accurately measuring the physicochemical properties of NMs in the *in vitro* setting is crucial. In this study, we proposed a physchem. score to obtain an indication of the level of details in the NM characterization in each study. This can lead to conflicting results about the link between material properties and cellular uptake. NMs can enter cells through various mechanisms, like endocytosis or passive diffusion. The dominant pathway can depend on the specific NM characteristics, adding another layer of complexity. In the majority of the studies, the physchem. characterization of the selected NM was assessed taking into consideration these parameters. However, the variability in both NM properties and assay methodologies poses significant challenge for comprehensive study comparisons. Finally, the dynamic extracellular environment of the gut significantly influences NM interactions with the cell membrane and subsequent uptake. Our literature research confirmed that while reporting physchem characterization of NMs has become more common in recent studies, the minimum essential information, such as NM characterization in digestive fluids or cellular media, is still not consistently reported using standardized methods. This hinders the establishment of reliable causal relationships between physchem properties and biological effects. Several studies suggest that the physicochemical characteristics of the NM, such as size, surface charge, and coating materials may have a role in promoting or hampering the cellular endocytic uptake (39,40). For instance, smaller size and surface modifications enhance silica NM uptake, as shown by increased uptake of smaller particles (30-130 nm) compared to larger ones (200 nm) (41). Another study investigated the uptake and translocation of Polyacrylic Acid (PAA)-coated Ag NMs and FeO NMs. While both types were taken up by cells, only FeO particles could cross the in vitro barrier. This suggests the core material of the NM, not the coating, plays a crucial role in translocation (42). Furthermore, ingested NMs might undergo complex transformations with the dynamic physicochemical environment of the gastrointestinal tract, which significantly influences their interaction and uptake by the biological system. Consequently, accurately modeling the food matrix effect and the unique GIT exposure conditions is essential for comprehensive NM bio-interaction studies. (38,43).

#### Systems

The differentiated Caco2 model and the triculture system Caco-2/HT29-MTX or Caco-2-TC7/HT29-MTX represent the traditional co-culture model to investigate the intestinal barrier functionality *in vitro* (37). Discrepancies between studies can also be attributed to variations in the timing and method of NM administration within the transwell system, specifically whether the NM was added to the apical or basolateral compartment. The commonly used Caco-2 cell line, primarily employed for assessing cell viability, contributes to inconsistencies in observed biological effects and dose-response relationships. This is because different cell types can exhibit varying sensitivities to NMs. For instance, Antonello et al. tested also primary non-transformed intestinal epithelial cells HCoEpiC and the colon cancer derived HCT116 cell line in addition to proliferative Caco2, and observed cytotoxic effect of FeO NMs in all cell lines with different grade of sensitivity (primary non-transformed cells > Caco-2 and HCT116 cells) (38).

## 4. Discussion

The European Food Safety Authority (EFSA) acknowledges the importance of Adverse Outcome Pathways (AOPs) in risk assessment. Their guidance document on NMs emphasizes the role of AOPs in providing a deeper mechanistic understanding of potential human health impacts. Furthermore, using AOPs alongside Integrated Approaches to Testing and Assessment (IATA) can streamline future testing of NMs in food facilitating the weight-of-evidence approach and minimizing further animal studies. However, while the AOP-Wiki is a valuable resource for how NMs can impact organs like lungs, liver, and the reproductive system, information on their effects on the intestine is scarce. This causes a knowledge gap and creates challenges in understanding the relevance of *in vitro* assays needed for the establishment of IATAs as described in Step 2 of the nano-guidance document. Scientific evidence, documented in the literature, demonstrates that internalized NM can disrupt the normal function of intestinal epithelial cells and paves the way for suggesting additional AOPs to elucidate the potential mechanisms by which NMs used in the food sector might exert adverse effects on the gut. By building upon existing AOPs with cytotoxicity as a key event, we developed a novel AOP, AOP530, and incorporated it into the AOP-Wiki. This AOP specifically addresses the potential disruption of the intestinal barrier by food-borne NMs. The presented AOP hypothesized that the endocytosis of specific NMs acts as the MIE for intestinal barrier dysfunction. A critical KE within this pathway is the cytotoxicity observed in enterocytes and goblet cells. These cell types contribute to the establishment of the barrier by expressing TJs relevant for the barrier function. However, it is crucial to acknowledge the inherent complexity of cellular responses to NMs, which often exhibit a non-linear and interconnected nature, as opposed to a strictly sequential order. By reviewing scientific literature, we investigated the documented biological effects of five NMs on the intestinal epithelium often used in the food sector.

### 4.1 Endocytic uptake as molecular initiating event (MIE)

Endocytosis is of particular interest in nanotoxicology due to its responsibility for the Trojan horse effect, in which NM facilitates the internalization of contaminants or serves as precursors, releasing breakdown products that would otherwise exhibit limited cellular permeability. This holds true for several metal oxide NMs. Upon internalization, these NMs can undergo degradation, leading to the uncontrolled release of metal ions and bypassing the regulatory mechanisms associated with metal transporters. In our study, we could reveal that relevant NMs for the food sector can be taken up by intestinal cells *in vitro*.

### 4.2 Endocytosis of NM can lead to lysosomal disruption (KER1775)

In our study, three of the analyzed NMs (ZnO, CuO and Ag NM) are known to release ions after accumulating in the acidic environment of lysosomal vesicles. The ionic forms Zn2+, Fe2+, Ag+ or Cu2+ can disrupt the stability of the lysosomal membrane through interactions with the phospholipid bilayer altering its structure or suppressing enzyme activities involved in maintaining the lysosomal integrity (72). The resulting leakage of digestive enzymes can further damage the cellular components and trigger cell death. In addition, an overload of positive ions in lysosomes also triggers anionic and water influx ultimately leading to a disruption of the lysosome. In addition, the NM itself can exhibit cationic characteristic depending on the surface of the nanoparticles and the specific environments. Such characteristics are described for FeO NMs and SiO_2_ NMs that can lead to an influx of anions and water (proton sponge effect). Lysosomes can swell and lead to the disruption of the lysosomal membrane (73,74) leading to lysosomal dysfunction. However, none of the investigated studies measured the disruption of lysosomal structures in enterocytes or goblet cells and preventing us to establish a direct linkage between endocytosis and lysosomal disruption, which is described in KER1775 for other NMs.

### 4.3 Lysosomal disruption leads to mitochondrial dysfunction (KER993)

Our hypothesis proposes that lysosomal disruption triggers the release of lysosomal proteases, consequently inducing mitochondrial dysfunction as described in KER993. Mitochondrial dysfunction encompasses a broad spectrum of alterations in both mitochondrial structure and function. NMs derived ions interfere with the redox cycling in mitochondria leading to an elevated ROS production. But also physical interactions of the entire NM with mitochondria and reactive groups on the NP surface can trigger ROS generation. This aspect is confirmed by several studies related to Ag NMs (39,40,78,45,63–65,70,75–77). As the concentration of NMs/ions increases within a cell (e.g. through accumulation after chronic or high exposure), the likelihood of interactions with cellular components, particularly mitochondria, also increases. Moreover, the production of ROS can initiate a cascade of additional cellular events with an impact on the intestinal barrier including inflammatory responses (18,79). The study of Xu and colleagues described an increased ROS production, pro-inflammatory cytokines, and cytotoxicity after treatment with smaller SiO_2_ nanoparticles affecting signal transduction pathways (80). Pathways such as RhoA/ROCK leading to the disassembly of TJ, inflammatory signaling cascades and cytoskeleton can compromise cellular processes crucial for a healthy barrier (45,78,80,81). Such impairments would not automatically lead to a cell death but can be of relevance for the intestinal barrier (57).

### 4.4 Mitochondrial dysfunctions lead to Cytotoxicity (KER363)

The causal relationship between mitochondrial dysfunction and cell death/injury is well captured within KER363 in the OECD-endorsed AOP48 and AOP144. ROS plays a critical role in cell death in particular at higher concentrations. A lower level of ROS production can act as a signaling molecule, even enhancing cellular stress resistance. However, when ROS production overwhelms the cell’s antioxidant defenses, a state of oxidative stress ensues promoting cell death. Physicochemical properties, the susceptibility of different cell types as well as exposure times are other parameters that can influence different cytotoxicity results. Increased lipid vacuoles and changes in morphologic ultrastructure like villi and organelles dimension has been also observed in NM treated cells (60). Moreover, altered expression of cytoskeleton and lipid-metabolism related proteins has been also found in *in vivo* studies (82,83). Passive lipid uptake could be progressively normalized through a decrease in intestinal adsorptive area because of changes in microvilli length, width and density of enterocytes. All these changes can contribute to membrane instability, lipid accumulation, oxidative stress, inflammation which ultimately lead to cell death machinery activation and intestinal epithelium homeostasis dysregulation (84,85). The balance between proliferation and apoptosis of intestinal cells is dependent on the microenvironment and stress factors; the defect in balance is strongly connected with several intestinal diseases and GI injuries (86,87).

### 4.5 Intestinal cell death leads to intestinal barrier disruption (KE3197)

The intestinal barrier is a multilayer system. Alteration of one or more layers of the intestinal barrier leads to increased intestinal permeability and barrier function reduction (88). Some studies have reported the role of TJs in regulating epithelial cell proliferation. For instance, overexpression of claudin-2 in human colon cells increased proliferation in vitro and accelerated tumor growth in vivo (89,90). Moreover, it is known that pro-inflammatory cytokines such as TNF and IL1β or LPS can induce barrier loss by internalization of occludin (91,92). Administration of recombinant IL-13 in mice increases caudin-2 expression and augments intestinal paracellular cation permeability (93). The causal linkage between damaged/dying enterocytes and goblet cells leads to increased paracellular permeability due to disruption of the epithelial monolayer and regulate mucus secretion and thickness, respectively, well described in literature (88,94) is now captured within KER3197 in the AOP-Wiki.

Of course, other pathophysiologic events leading to barrier disruption can occur in the gut after NM ingestion. For instance, some food additives have been shown to induce microbiota composition alterations. These microbiota alterations have been associated with a reduced mucus layer thickness (95) and an increased gut penetrability (96) linked with intestinal inflammation and metabolic alterations of the intestinal barrier (94).

## 5. Conclusions

This study proposes a novel AOP linking NM uptake to intestinal barrier disruption. While the proposed mechanism is biologically plausible, the available evidence, primarily derived from studies on ZnO, CuO, FeO, SiO_2_ and Ag NMs, offers limited support. The observed variability in study outcomes can be attributed to the heterogeneity in NM properties, biological systems, treatment type, and dose. To strengthen the AOP, further research is required, including systematic investigations of the proposed Key Event Relationship (KER) using well-characterized stressors. Identifying a prototypical stressor would also enhance the AOP’s utility for quantitative assessments. It is essential to consider that other AOPs, such as those involving inflammation, may also contribute to intestinal barrier impairment induced by NMs. A comprehensive understanding of NM toxicity requires the integration of multiple AOPs. Despite these limitations, the proposed AOP provides a valuable framework for understanding the potential toxicity of existing and emerging food NMs, guiding future research and risk assessment efforts.

## Supporting information

Tables and boxes

