## Supplementary material for "An Adverse Outcome Pathway for Food Nanomaterial-induced Intestinal Barrier Disruption": Tables and boxes

**Glossary**
**D** Differentiated Caco2
**PL** Proliferative Caco2
**PT:** Cell culture in plate
**T:** Cell Culture in transwell system
**G:** Cell culture on glass slide
**AP:** apical treatment in transwell
**BL:** basolateral treatment in transwell
**H&E:** hematoxilin-eosin staining
**AB-PAS:** Alcian blue-PAS staining
**LY:** luciferase yellow
**TEER:** transepithelial resistance
**IF:** immunofluorescence
**Papp:** Apparent permeability
**IHC**: immunohistochemistry
**LM**: light microscopy
**TEM**: Trasmission electron microscopy
**SEM**: scanning electron microscopy
**IBM** ion beam microscopy
**AFM** Atomic force microscopy
**CPE**: cloud point extraction
**CLSM** confocal laser scanning microscopy
**CBMN** cytokinesis-block micronucleus
**CTB** CellTiter-Blue
**FCMN** flow cytometric micronucleus
**ICP-MS**
**ICP-AES**
**ICP-OES**: Inductive Coupled Plasma Optical Emission Spectrometry
**AAS**:Atomic absorption spectrometry
**NTA:**.Nanoparticle Track Analysis
**IBCA** Impedance-based cellular assays
**GSEA** Gene set enrichment analysis
**WB** Western blot
**EDS** Energy Dispersive X-ray Spectroscopic

Tables and boxes

**Table 1.** AOPs reporting NMs as stressors in the AOP-Wiki.

| stressor | AOP ID | MIE | AO | status |
|---|---|---|---|---|
| NPs | 144 | Endocytotic lysosomal uptake | Liver fibrosis | **OECD under review (1.47)** |
| carbon nanotubes and carbon nanofibres | 173 | Substance interaction with the lung resident cell membrane components | Pulmonary fibrosis | **OECD approved (1.32)** |
| insoluble nanosized particles | 237 | Substance interaction with the lung resident cell membrane components | Atherosclerosis | **Under development (1.55)** |
| high aspect ratio material | 303 | Frustrated phagocytosis | Lung cancer | **Under development (1.86)** |
| "wide range of nanomaterials" | 302 | Inhibition of lung surfactant function | Decreased lung function | **Under development (1.87)** |
| nanoparticles, SARS-CoV-2 | 319 | Induced dysregulation of ACE2 | lung fibrosis | **Under development (1.96)** |
| silica NPs | 481 | Increase, Reactive Oxygen Species | respiratory dysfunction | not OECD |
| carbon nanotubes | 241 | Activation, Latent Transforming Growth Factor Beta 1 | Pulmonary fibrosis | not OECD |
| nanomaterials | 451 | Substance interaction with lung resident cell membrane components | lung cancer | not OECD |
| carbon nanotubes | 409 | Frustrated phagocytosis | increase, mesotheliomas | not OECD |
| nanomaterials, SARS-CoV-2 | 392 | Fibrinolysis, decrease | hyperinflammation | **Under development (1.96)** |
| silver NPs | 207 | Activation, NADPH oxidase | Reproductive failure | not OECD |
| titanium dioxide NPs | 208 | ? | Reproductive failure | not OECD |
| silica NPs | 209 | ? | Hepatotoxicity | not OECD |
| graphene oxide NPs | 210 | ? | Reproductive failure | not OECD |

**Table 2.** Evidence from published literature supporting AOP530 for ZnO NM.

| NM | PC | System | KE1539 Endocytotic lysosomal uptake | | KE898 Lysosomal disruption | | KE177 Mitochondrial dysfunction | | KE55 Cell death/injury | | KE1931 Intestinal barrier disruption | | Ref. |
|---|---|---|---|---|---|---|---|---|---|---|---|---|---|
| | | | +/-/= | Assay | +/-/= | Assay | +/-/= | Assay | +/-/= | Assay | +/-/= | Assay | |
| ZnONP (544906, <50um 67745, <100um Sigma-Aldrich) | 3 | Caco-2 DT | + | ICP-MS (614uM) | | | = | MTT | = | MTT | = = | TEER Papp (614uM) | (1) |
| | | HT29-MTX DT | + | ICP-MS (614uM) | | | = | MTT | = | MTT Alcian blue staining | - = | TEER Papp (307uM) | |
| ZnONP (544906 Sigma-Aldrich) | 3 | Caco-2 DT | | | | | | | | | + | TEER (100ug/ml) | (2) |
| ZnONP (<100nm Sigma Aldrich) | 3 | Caco-2 P P | + | ICP-AES | | | | | | | | | (3) |
| Digested ZnONP (544906, <50um 67745, <100um Sigma-Aldrich) | 3 | Caco-2 DT | + | ICP-MS (307uM) | | | = | MMP | = | MTT, TEM | = | TEER Papp (614uM) | (4) |
| | | Caco-2 + HT-29 DT | + | ICP-MS (307uM) | | | = | MMP | = | MTT, TEM | = | TEER Papp (614uM) | |
| Intragastric ZnONP (<50nm Sigma Aldrich) | 3 | 30 days, mouse | | | | | | | + | H&E | + | Histology (mucosa) RT qPCR (TJ) | (5) |
| Digested ZnONP (<100nm) | 0.5 | Caco-2 D | | | | | + | TEM | + | MTT, LDH 0.523uM | | | (6) |
| Digested ZnONP | 3 | Caco-2 P, DT | | | | | | | - | Calcein AM/PI (0..097mg/ml) | - | TEER IF (TJ) | (7) |
| digested ZnO NP (nanoscale material <10nm) | 0.5 | Caco-2 PP | | | | | + | WST-1 (20ug/cm2 | + | LDH WST-1 | | | (8) |
| ZnO NP (nanoscale material <10nm) | 3 | Caco-2 D | + | ICP-OES | | | + | WST-1 (20ug/cm2 | + | WST-1 | | | (9) |
| NM110 | 3 | Caco-2 transwell/ gut-on-chip | | | | | + | WST-1 (50ug/ml) | + | WST-1 (50ug/ml) | | | (10) |
| ZnO NPs (Sumitomo) | 3 | Caco-2 PP | + | ICP-AES | | | + | WST-1 (10ug/ml) | + | WST-1 (10ug/ml) | | | (11) |
| ZnO + vit C + CPP | 3 | GES-1 | | | | | + | MMP | + | LDH | | | (12) |
| ZnO NPs (American Elements) | 2.5 | INT-407 porliferating | + | ICP-AES | | | | | + | LDH (50ug/ml) | | | (13) |
| ZnO NPs (Aladdin 30nm) | 0.5 | In vivo mouse colon | | | | | | | | | + | Histology (mucosa) RT qPCR (TJ) | (14) |
| ZnO /Zn2+ (Merck Corp) | 3 | In food, 270d mouse jejunum | = | TEM | = | TEM | | | = | histology | + | Histology (mucosa) RT qPCR (TJ) | (15) |

Tables and boxes

**Table 3.** Evidence from published literature supporting AOP530 for CuO NM.

| NM | PC | System | **KE1539** Endocytotic lysosomal uptake | | **KE898** Lysosomal disruption | | **KE177** Mitochondrial dysfunction | | **KE55** Cell death/injury | | **KE1931** Intestinal barrier disruption | | Ref |
|---|---|---|---|---|---|---|---|---|---|---|---|---|---|
| | | | +/-/= | Assay | +/-/= | Assay | +/-/= | Assay | +/-/= | Assay | +/-/= | Assay | |
| CuO NM (10nm, Plasma Chem, GmbH) | 3 | Caco2/ RajiB, TD AP<br>Caco2/ RajiB-MTX TD AP | + | ICP-OES | | | | | = | SEM<br>IF nuclei<br>Romanowsky staining | +<br>+<br>+ | TEER<br>IF (ZO1)<br>Papp | (16) |
| CuO (c)ommercial (Sigma),(e)thanol, and (w)ater | 2 | Caco2 TD AP or BL | | | | | | | | | + | TEER CuOc (10 µg/mL, BL); CuOe, CuOw (100µg/mL, AP) | (17) |
| CuO (<50nm, Sigma) | 3 | EpiIntestinal™ (SMI-100) | | | | | + | MTT | + | MTT (40µg/mL) | | | (18) |
| | | Rat IEC-6 | | | | | + | Mitotracker (0.08µg/mL) | + | MTS 4µg/mL | | | |
| CuO (10nm, Plasma Chem, Gmbh, Berlin) | 3 | Caco2 PL, TD AP | + | ICP-OES | | | + | Alamar blue (2.44 µg/cm²) | + | Alamar blue (2.44 µg/cm²) | +<br>+<br>+ | TEER<br>IF (ZO1)<br>IF (DAPI) | (19) |
| CuO in lettuce | 0.5 | Caco2 TD, AP 2h | + | TEM | + | TEM | + | TEM | +<br>+ | LDH<br>MTT | | | (20) |

**Table 4.** Evidence from published literature supporting AOP530 for FeO NMs.

| NM | PC | System | KE1539 Endocytotic lysosomal uptake +/-/= | KE1539 Assay | KE898 Lysosomal disruption +/-/= | KE898 Assay | KE177 Mitochondrial dysfunction +/-/= | KE177 Assay | KE55 Cell death/injury +/-/= | KE55 Assay | KE1931 Intestinal barrier disruption +/-/= | KE1931 Assay | Ref |
|---|---|---|---|---|---|---|---|---|---|---|---|---|---|
| VENGES Fe2O3 Digested (300nm) | 3 | Caco-2, HT29, RajiB TD - AP | + | ICP-MS 2/4h | | | | | = | Live/Dead LDH | = | TEER | (21) |
| PLGA-PEG Fe3O4 Digested (100nm, Colorobbia Cosulting , Italy) | 3 | Caco2 PT/PL | | | | | + | WST-1 50μg/ml | + | WST-1 50μg/ml | | | (22) |
| | | HCT116 | | | | | = | WST-1 | = | WST-1 | | | |
| | | HCoEpiC | | | | | + | WST-1 20μg/m | + | WST-1 20μg/mL | | | |
| | | Caco2 TD AP | = | NTA Ferrozine | | | = | WST-1 | = | WST-1 | =<br>= | TEER<br>LY | |
| | | | | | | | | | | | - | qRT-PCR (TJP1, OCLN and CDN5) | |
| E172 2 Yellow FeO(OH), 2 Red Fe2O3, 1 Orange Fe2O3+FeO (OH) and 2 Black Fe3O4 (<170 nm) Digested | 3 | Caco2 PT/PL | | | | | = | MTT 100 μg/mL | =<br>= | MTT 100 μg/mL<br>IBCA Red 1, 2 - Orange | | | (23) |
| | | | | | | | | | + | IBCA Yellow 1/2 Black 1/2 >5 μg Fe/mL | | | |
| | | Caco2 TD, AP | + | AAS | | | | | = | IBCA | | | |
| Amino PVA USPIO (polyvinylialchol/polyinylamine amino) (40nm) | 3 | Caco-2 and HT29 spheroids. | + | Prussian blue, nuclear Red | | | | | | | | | (24) |
| | | Caco2 HT29 PL/PT | +<br>- | Prussian blue, nuclear Red TEM (21μg/mL) | | | = | MTT | = | MTT | | | |
| | | Caco2 T/D | | | | | | | | | = | LY | |
| | | Caco2/HT29 T/D - AP | | | | | | | | | = | LY | |
| PAA coated Fe3O4 NM (200nm, Chemichell) | 3 | Caco2 T/D | + | AAS TEM IBM 14μg/mL | | | | | = | Cell Titer Blue | | | (25) |
| | | Caco2/HT29/RajiB T/D - AP | + | AAS | | | | | | | = | TEER Papp 200μg/mL | |
| FeO digested (698nm, China n/a) | 2 | mice,150 mg/kg/d, 30 days Intragrastric. | | | | | | | + | H&E | | | (26) |
| | | Caco-2 T/D 100μg/mL AP | | | | | | | | | + | Papp 100μg/ml | |

Tables and boxes

**Table 5.** Evidence from published literature supporting AOP530 for SiO2 NMs.

| NM | PC | System | **KE1539** Endocytotic lysosomal uptake | | **KE898** Lysosomal disruption | | **KE177** Mitochondrial dysfunction | | **KE55** Cell death/injury | | **KE1931** Intestinal barrier disruption | | Ref. |
|---|---|---|---|---|---|---|---|---|---|---|---|---|---|
| | | | +/-/= | Assay | +/-/= | Assay | +/-/= | Assay | +/-/= | Assay | +/-/= | Assay | |
| Sicastar - Red (micromod) | 3 | ISO-HAS-1/ Caco-2 DT | | | | | | | | | = | TEER NaFlu | (27) |
| SiO2 nanobeads (HiQ-Nano srl) | 3 | Caco2-HT29MTX-Raji DT | + | CLSM | | | | | | | + | TEER IF (TJ) 100 μg/ml | (28) |
| precipitated and fumed SiO2 | 3 | Caco2 PT-D Caco2/Raji B - T PL - AP | + | CPE 2h | | | | | | | + | Papp 6h | (29) |
| SiO2 | 3 | C57BL/6J mice, 3g/kg/day, 28 days | | | | | | | + + + | H&E AB-PAS IHC | + | IF TJ | (30) |
| Kaolinite (Argiletz, France) | 0 | Male Wistar rats, 3.4g/day in food 28 days | + | ESEM/EDX | | | | | = | LM TEM SEM | = | proteomic | (31) |
| Kaolinite (Argiletz, France) | 0 | Male Wistar rats | = | TEM | | | | | = | TEM | | | (32) |
| SAS Food-grade SiO2 nanoparticles (E551) dissolution study 24h | 2.5 | Caco-2 BBe1, DG, AP | | | | | | | | | + | SEM 1 ug/mL 24h | (33) |
| SAS D90 (DMSNs, Spherical, 90, 130 nm) | 3 | Caco-2/HT29-MTX-E12 PT, D | + | IF | | | | | = | AFM IF | = | TEER IF | (34) |
| SAS (10-200 nm) | 3 | Caco-2 Caco-2/HT29-MTX TD - PT AP, BL | + | CLSM | | | | | = | MTT LDH ROS | + | TEER CLSM Papp ZO-1 (NP30) | (35) |
| SAS | 3 | DLD-1 SW480 NCM 460 PL, PT | + | CLSM | | | | | = | MTT, Annexin V/PI ROS | = | TEER CLSM | (36) |
| SiO2 FG-NP, FG-MP, NFG-NP | 2.5 | Caco-2 PL, PT | | | | | | | = | MTT, LC20 (62.8 ppm). | + | GSEA | (37) |
| SiO2 E 551 | 2.5 | HT29-MTX-E12 PL, PT | + | FACS IF | | | | | = | MTS FACS SCGE | | | (38) |
| SAS | 3 | Male Sprague | + | HP | | | | | = | SCGE HP | | | (39) |

| | | | | | | | | | | |
|---|---|---|---|---|---|---|---|---|---|---|
| | | Dawley rats | | | | | | | | MDA |

**Table 6.** Evidence from published literature supporting AOP530 for AgNMs.

| **NM/provider/size pristine/digestion** | **PC** | **Cell Model** | **KE1539** Endocytotic lysosomal uptake | | **KE898** Lysosomal disruption | | **KE177** Mitochondrial dysfunction | | **KE55** Cell death/injury | | **KE1931** Intestinal barrier disruption | | **Ref** |
|---|---|---|---|---|---|---|---|---|---|---|---|---|---|
| | | | +/-/= | Assay | +/-/= | Assay | +/-/= | Assay | +/-/= | Assay | +/-/= | Assay | |
| NM300K (Fraunhofer IME, Germany) 7.74 ± 2.48 nm TEM | 3 | Caco-2<br>DT<br>PL<br>AP | + | ICPMS<br>TEM | | | | | + | comet assays<br>50 µg/mL | = | TEER<br>Papp<br>50µg/mL<br>qRT-PCR (TJ) | (40) |
| PAA-coated Ag | 2 | Caco-2, HT29-MTX-E12 Raji-B<br>DT, AP | + | AAS<br>TEM SEM | | | | | = | cell titer blue<br>20 µg/mL | = | TEER<br>Papp<br>20 µg/mL | (41) |
| AgNP<br>4.84 ± 2.1 nm | 2.5 | Caco-2<br>HT29<br>DT, AP | + | CLSM | | | | | | | + | TEER<br>Papp<br>RTq-PCR (TJ)<br>100µg/mL | (42) |
| Ag NPs and AgNO3<br>20-200 nm | 2 | Caco-2/TC7<br>HT29-MTX<br>DT PL AP | + | CLSM | | | | | = | Alamar blue<br>100µg/mL | | | (43) |
| Ag NP<br>35 nm | 2 | C3a<br>Caco-2<br>DT<br>PL<br>AP | + | CLSM | | | | | + | LDH IC50<br>50 µg/mL | | | (44) |
| AgNP<br>20,34,61,113 nm | 3 | Caco-2-RajiB<br>DT, AP | | | | | | | | | = | TEER<br>Papp<br>25 µg/mL | (45) |
| AgPURE Rent a Scientist GmbH, TEM 7.02 ± 0.68 nm/Y | 3 | Caco-2<br>DT, PL | | | | | | | + | CellTiter Blue<br>DAPI<br>15µg/mL | + | electrodes | (46) |
| Ag from mussels<br>23nm | 0 | Caco-2<br>DT, AP | + | TEM | | | | | | | = | TEER<br>LY<br>0.0406 to 0.265 µg/g | (47) |
| NM300K<br>20nm | 1.5 | Caco-2<br>DT, AP | | | | | | | = | LDH<br>15 µg/mL<br>(3h) | | | (48) |
| NM300<br>Ras GmbH, 20nm | 2 | Caco-2<br>HT29MTX DT, AP | + | ICP MS<br>40ug/mL | | | | | | | | | (49) |
| poly (acrylic acid)-coated AgNM<br>3.2±0.1 nm/Y | 2 | Caco-2<br>DT<br>AP | + | AAS | | | | | + | Cell titer blue<br>40ug/mL | = | TEER<br>Papp<br>20 to 100 µg/ml | (50) |
| AgPURE | 3 | Caco-2,<br>D, PT | + | AAS | = | Proteomics | + | Proteomic | + | NRU, MTT, CTB, DAPI | = | Proteomic<br>TEER | (51) |

| | | | | | | | | | | | | | |
|---|---|---|---|---|---|---|---|---|---|---|---|---|---|
| AgNO3 | | | + | AAS | = | Proteomics | + | Proteomic | + | NRU, MTT, CTB, DAPI | = | Proteomic TEER | |
| AgNO3 | 2.5 | Caco-2, PL, PT | + | TEM-EDX, ICP-MS | + | CLSM | + | ROS 5 µg/mL IPA | + | Cell counting Comet assay 5 µg/mL | | | (52) |
| AgNM | 3 | Caco-2, PL, PT | + | TEM-EDX ICP-MS | + | CLSM | + | ROS IPA | + | Comet Assay | | | |
| AgNM | 3 | Caco-2 D PT | + | CLSM TEM | | | | | = | LDH | | | (53) |
| Micro-Ag | 2.5 | Caco-2 DT AP | + | CLSM TEM | | | | | = | LDH | | | |
| Ag-PVP (Polyvinylpyrrolidone-capped Silver) | 3 | Caco-2 HT29-MTX-E12 THP-1, PL, PT T, AP BL | + | SEM | | | + | ROS | + | LDH WST-1 Comet Assay | = | TEER | (54) |
| Ag-NPs | 2.5 | Caco-2, SW480 PL, PT | + | TEM | | | = | ROS (DHE) | + | MTT (100 mg/L) | | | (55) |
| Ag-NPs | 2.5 | NCM460 HCT116 PT, PL | + | TEM | = | Protein | + | ROS WB | + | MTT WB LDH RT-qPCR | = | microscopy | (56) |
| (Polyethyleneimine) PEI-AgNP 4 nm AgNP 19 nm AgNP | 3 2.5 | Caco-2 PL, PT | + | TEM FTIR | | | + | ROS NO | + | MTT Annexin V/PI (4.5µg/ml), LDH | | | (57) |
| Ag Nanoparticles Digested NP | 2.5 | Caco-2 PL, PT | + | TEM EDS | | | | | + | MTT, TEM | | | (58) |
| Pristine AgNPs, Digested AgNPs, Silver nitrate (AgNO3) | 3/ 3 | Caco-2 / HT29-MTX D, PT AP, BL | + | CLSM spICP-MS | = | CSLM | | | + | WST-1 CLSM ICP-MS, spICP-MS | = | TEER, LY FITC-D | (59) |
| Silver nanoparticles (AgPURE) | 2.5 | Caco-2, PL, PT, D | + | TEM-EDX | | | + | Proteomic | = | Cell Titer Blue | + | qRT-PCR | (60) |
| Peptide-coated AgNM | 2 | Caco-2, PL, PT, D | + | TEM-EDX | | | + | ROS (DCFH-DA) | + | CTB Annexin-V/7AAD LDH DAPI | + | IBCA | (61) |
| AgNM Ag-AuNM | 2.5 | HuTu-80, PL, PT | + | TEM | | | + | ROS | + | IBCA | | | (62) |
| AgNM AgNO3 | 3 | Caco-2 THP-1 D, T, AP, BL | + | TEM | | | | | + | LDH DAPI | + | TEER | (63) |
| AgNM +/- coatings | 2.5 | Caco-2, PL, PT | + | TEM | | | + | ROS | + | LDH Annexin- V FITC/PI DAPI | | | (64) |
| AgNM | 2.5 | Caco-2, D, T, AP, BL, | + | TEM ICP-MS | | | | | + | TEER LDH | + | TEER | (65) |
| AgNM | 2 | Caco-2, PL, PT | + | TEM ICP-MS | + | MN Acridine orange | | | + | Trypan blue | | | (66) |

Tables and boxes

| | | | | | | | | | | | | | |
|---|---|---|---|---|---|---|---|---|---|---|---|---|---|
| | | | | | | | | | | Alamar blue | | | |
| AgNM (30nm) digested | 3 | Caco-2 PL, P | + | TEM ICP-MS | + | Proteomic | + | Proteomic | + | Proteomic | + | Proteomic | (67) |
| AgNM (20, 50nm) | 2/1.5 | Caco-2 PL, PT | + | TEM, ICP-MS | + | FCM | | | + | Alamar Blue, Trypan blue | | | (68) |
| AgNM (20, 50nm) | 2/1.5 | Caco-2 PL, PT | + | TEM, ICP-MS | + | CBM | | | + | Trypan blue | | | (69) |
| AgNM (14nm) | 3 | Caco-2, D, PT | + | TEM, ICP-MS | + | 2-DE, MALDI-TOF MS | + | Proteomics IPA | + | CTB DAPI | | | (70) |

**Table 7.** Evidence from published literature supporting AOP530 for nanocellulose.

| NM | PC | Cell Model | KE1539 Endocytotic lysosomal uptake | | KE898 Lysosomal disruption | | KE177 Mitochondrial dysfunction | | KE55 Cell death/injury | | KE1931 Intestinal barrier disruption | | Ref |
|---|---|---|---|---|---|---|---|---|---|---|---|---|---|
| | | | +/-/= | Assay | +/-/= | Assay | +/-/= | Assay | +/-/= | Assay | +/-/= | Assay | |
| CNC-140×20 nm; CNC-250× 25 nm; CNC-700× 25nm; PCNC-540x35 nm; CNF-50nm; CNF-80nm; TCNF-250×25nm synthetized | 3 | Caco2 TD | | | | | + | MTS CNC-250 CNF-80 (50µg/mL) | + | LDH CNC-250 (50µg/mL) | = = | Papp IF TJ(ZO1) CNF-50nm CNF-80nm | (71) |
| CNF/TiO2 (Nanostructured & Amorphous Materials Inc.) | 2 | Caco2, PL, PT | | | | | = | MTT 1000µg/ml | = | MTT 1000µg/ml | | | (72) |
| CNF (from wood pulp, University of Maine, 28 nm) | 0.5 | Caco2, FHC PL, PT | | | | | = | MTT WST-8 1000µg/ml | = | MTT WST-8 1000µg/ml | | | (73) |
| CNC (University of Maine, 5−20 nm width, 150−200 nm length) | 1 | Caco2 PT, D | | | | | = | MTT 10mg/mL | = | MTT 10mg/mL | = | Papp | (74) |
| CNC type I and II, (obtained from microcrystalline NC, Sigma, 5−10 nm width, 200−300 nm length) | 3 | Caco2/ TC7 PL and PD | | | | | = | MTT 5ng/µL | = | MTT 5ng/µL | | | (31) |

Tables and boxes

**Table 8.** Summary of weight-of-evidence (WoE) evaluation of key event relationships (KERs) in the AOP.

| KER ID | Upstream KE | Downstream KE | Biological Plausibility | Empirical Evidence | Overall WoE | Quantitative Understanding |
|---|---|---|---|---|---|---|
| 1775 | Endocytic lysosomal uptake | Lysosomal disruption | High | Low | Low | Low |
| 993 | Lysosomal disruption | Mitochondrial dysfunction | High | Low | Low | Low |
| 363 | Mitochondrial dysfunction | Cell death/injury | High | Moderate | Moderate | Low |
| 3197 | Cell death/injury | Intestinal barrier, disruption | High | Moderate | Moderate | Low |

**Box 1.**

- Transmission electron microscopy (**TEM**) is appropriate for visualizing NMs inside cells, since light microscopy fails to resolve them at a single particle level (Brandenberger et al., 2010; Villamil Giraldo et al., 2016).
- Confocal laser scanning microscopy (**CLSM**) combines high-resolution optical imaging with depth selectivity which allows optical sectioning (Harush-Frenkel et al., 2006; Zhang and Monteiro-Riviere, 2009).
- Inductively coupled plasma mass spectrometry (**ICPMS**) is a type of mass spectrometry capable of detecting metals and several non-metals at very low concentrations (Ng et al., 2015).
- Inductively coupled plasma optical emission spectrometry (**ICP-OES**) enables to detect the presence of elements based on their emission of light when excited by plasma energy
- Fluorescence-activated cell sorter (**FACS**) is a specialized type of flow cytometry for sorting a heterogeneous mixture of biological based upon the specific light scattering and fluorescent characteristics of each cell (Harush-Frenkel et al., 2006; Zhang and Monteiro-Riviere, 2009).
- Atomic absorption spectroscopy (**AAS**) is a widely used technique for the quantitative determination of elements in a variety of samples by measuring the absorption of specific wavelengths of light by the element of interest.
- EDS is used in conjunction with scanning electron microscopy (**SEM**) to analyze the elemental composition of materials.

**Box 2.**

- **Lysotracker green** (200 nM) is regularly used to assess lysosomal acidification (Anguissola et al., 2014, Wang et al., 2013; Kroemer and Jäättelä, 2005).
- Changes in morphology can be observed by using **acridine orange**, a weak base that accumulates in the acidic compartment of the cell mainly composed of lysosomes (Li et al., 2000; Reiners et al., 2002; Kroemer and Jäättelä, 2005). This is followed by flow cytometry (Zhao et al., 2001), static cytofluometry or flow cytofluometry (Antunes et al., 2001).
- Lysosomal membrane **permeabilization** can be visualized by immunostaining of lysosomal enzymes such as cathepsin B (Boya et al. 2003a). More specific staining can be achieved by **staining with antibodies** against lysosomal membrane proteins (Kroemer and Jäättelä, 2005).
- **Proteomics** analysis can offer a comprehensive view of the changes in protein expression, or post-translational modifications associated with lysosome disruption. By identifying alterations in protein profiles, proteomics can provide valuable insights into the molecular mechanisms involved in lysosomal dysfunction.
- **Confocal** microscopy enables high-resolution imaging of cellular structures, including lysosomes. By visualizing lysosomal morphology, localization, and dynamics, confocal microscopy can directly demonstrate alterations in lysosomes caused by disruption. It can also be used in conjunction with specific dyes or probes to track lysosome-related processes in real-time.

**Box 3**

- Mitochondrial membrane potential (**MMP**) measurement. **JC-1** staining is a common fluorescent dye-based assay used to assess mitochondrial membrane potential, which can be a direct indicator of mitochondrial health and function. In conjunction with fluorescence microscopy or flow cytometry, it is used to quantify changes in mitochondrial membrane potential
- Enzymatic activity of electron transport system via **MTT** assay is a colorimetric assay where NAD(P)H-dependent cellular oxidoreductase enzymes reflect the number of viable cells
- **ATP** content measurement via ATP assay used to signal the presence of metabolically active cells
- Cellular oxygen consumption

Tables and boxes

- Proteomic analysis analyzes changes in protein expression levels, post-translational modifications, and interactions within the mitochondria. proteins commonly associated with mitochondrial dysfunction include cytochrome c, heat shock proteins etc.

**Box 4**

- **WST-1**
- **MTT** assay is a colorimetric assay where NAD(P)H-dependent cellular oxidoreductase enzymes reflect the number of viable cells
- **ATP** assay used to signal the presence of metabolically active cells.
- **LDH** leakage assay: lactacte dehydrogenase (LDH) is a soluble cytoplasmic enzyme released outside the cell when the plasma membrane is damaged and detected with a tetrazolim salt.
- Propidium iodide (**PI**) is an intercaling and fluorescent molecule used to stain necrotic cells.
- **Neutral red** uptake is based on the ability of viable cells to incorporate and bind the supravital dye neutral red in lysosomes.
- Trypan blue assay
- **TUNEL** detects DNA fragmentation from apoptotic signaling cascades
- Caspase activity assays
- **Hoechst/DAPI** stain binds specifically to DNA molecules present in the cell nucleus if the cell membrane is not intact and the stain can penetrate. It is a fluorescent dye. When exposed to ultraviolet light, the dye bound to DNA emits blue light, making the nucleus easily visible under a fluorescence microscope.
- **Acridine Orange** visualizes nuclear changes and apoptotic body formation under a fluorescent microscope
- **Annexin V/PI Staining**
- Impedance-based cellular assays (**IBCA**). allow to non-invasively and instantaneously detect and monitor cell responses to chemical and biological agents. Small changes in the impedence of current flow in the cell culture substrate allow to determine the events of cell adhesion, spreading, growth, motility, and death (10.1098/rstb.2017.0226)
- Live/Dead staining is a mixture of two fluorescent dyes that differentially label live and dead cells.
- Hematoxilin and Eosin (**HE)** staining of tissue sections: the nuclei are stained purple, while the cytoplasmic components are pink)

**Box 5**

Paracellular permeability

- Transepithelial electrical resistance (**TEER**) measure barrier integrity of the cell layer.
- Monitoring the passage of fluorescent molecules **(FITC**-dextran, sodium fluorescein or Luciferase yellow **(LY),** provides the apparent permeability coefficient **(Papp)** after the using the formula Papp= (dQ/dt)/AC0 where dQ/dt is the trasport drug/NM per unit time (mg/s); A is the area of the transport membrane (cm2); and C0 is the initial concentration of sample solutions (μg/mL).
- Staining of tight junctions (**TJ**), such as ZO-1, occludin by immunofluorescence

Mucus secretion and thickness

- **Alcian blue** staining can visualize the mucus layer, structure and thickness
- **Histology** staining (HE) to assess intestinal mucosal thickness and number of goblet cells
